# Vireo: Bayesian demultiplexing of pooled single-cell RNA-seq data without genotype reference

**DOI:** 10.1101/598748

**Authors:** Yuanhua Huang, Davis J McCarthy, Oliver Stegle

## Abstract

The joint analysis of multiple samples using single-cell RNA-seq is a promising experimental design, offering both increased throughput while allowing to account for batch variation. To achieve multi-sample designs, genetic variants that segregate between the samples in the pool have been proposed as natural barcodes for cell demultiplexing. Existing demultiplexing strategies rely on access to complete genotype data from the pooled samples, which greatly limits the applicability of such methods, in particular when genetic variation is not the primary object of study. To address this, we here present Vireo, a computationally efficient Bayesian model to demultiplex single-cell data from pooled experimental designs. Uniquely, our model can be applied in settings when only partial or no genotype information is available. Using simulations based on synthetic mixtures and results on real data, we demonstrate the robustness of our model and illustrate the utility of multi-sample experimental designs for common expression analyses.

## Background

Single-cell RNA-seq (scRNA-seq) is a rapidly evolving technology. Robust protocols and reduced costs have fostered applications in biomedicine, for example to identify biomarkers in disease [1, 2], or to characterize the cellular response to treatment and other external stimuli [3, 4].

Across these use cases, pooled experimental designs that combine multiple samples in a single experiment have critical statistical advantages compared to the serial analysis of cells from multiple samples in independent experimental batches [5, 6]. In particular, pooled designs allow dissecting true inter-individual variation from experimental batch variation. Pooled designs whereby a large number of cells are processed in a joint fashion are facilitated by the uptake of droplet sequencing methods in particular, including Drop-seq [7] and the 10x Genomics Chromium platform [8], which can assay tens of thousands of cells in a single run.

The aforementioned advantages have motivated a series of barcoding strategies to demultiplex samples from pooled experiments. In addition to simplified experimental logistics and reduced batch variation, pooled designs can also facilitate the identification of double cells. Existing barcoding strategies include molecular labelling prior to analysis [9, 10, 11, 12] as well as exploiting natural genetic barcodes of germline variants that segregate between pooled individuals [13]. While molecular barcoding is in principle applicable to any study design, genetic barcoding is both elegant and can be seamlessly integrated in existing scRNA-seq workflows, without the need to introduce additional processing steps.

The multiplexed designs with genetic barcoding are particularly useful in biomedical research, where the analysis of larger cohorts of genetically distinct individuals is particularly relevant [14]. However, current methods for demultiplexing genetically barcoded pools, such as Demuxlet [13], require genotype reference data for the pooled samples. Using variant information extracted from the scRNA-seq reads, each cell is assigned to a sample in the pool based on its genetic distance to the known genotypic states in a predefined reference database. While there is a growing interest in multi-sample analyses to study the effect of genetic variation between individuals at single-cell level, e.g., [15, 16], the requirement to supply a genotype reference database is prohibitive for studies without a genetic focus *per se*. Consequently, the potential of pooled experimental designs is currently not fully realized.

To address this, we here present Vireo (Variational Inference for Reconstructing Ensemble Origins), a principled Bayesian method to demultiplex arbitrary pooled designs that combine genetically distinct individuals. Uniquely, Vireo models the genotypes of each individual as latent variables, which are inferred from the observed scRNA-seq reads. The model can also leverage partial genotype information, e.g. when genotype data are available for a subset of individuals, and hence can be applied to a wide range of experimental settings.

## Results and discussion

Vireo jointly assigns each cell to one of *K* individuals and estimates the genotypic state of these individuals at known polymorphic loci. The model takes a set of common genetic variants as input (for example derived from the 1000 Genomes Project [17]), which are genotyped in each cell based on the scRNA-seq read data. Despite the typically low coverage of single-cell RNA-seq experiments, this approach allows for genotyping on the order of 100 expressed variants per cell (e.g. using 3’ 10x Genomics data; approx. 50,000 reads per cell, Fig 1 and Methods). Combining information across cells, these sparse genotype data allow for reconstructing the underlying individuals that are represented in the pool, which in turn allows for assigning each cells to an individual (Fig. 1). Vireo also explicitly accounts for doublets (two or more cells processed as a “single cell” in the assay) by considering cells that appear to be assigned to a combination of individuals. Finally, the model estimates the most likely number of pooled individuals, a feature that is useful if some of the pooled samples drop out for experimental reasons, and the method can incorporate partial genotype data that are available for a subset of the pooled samples.

**Figure 1.**
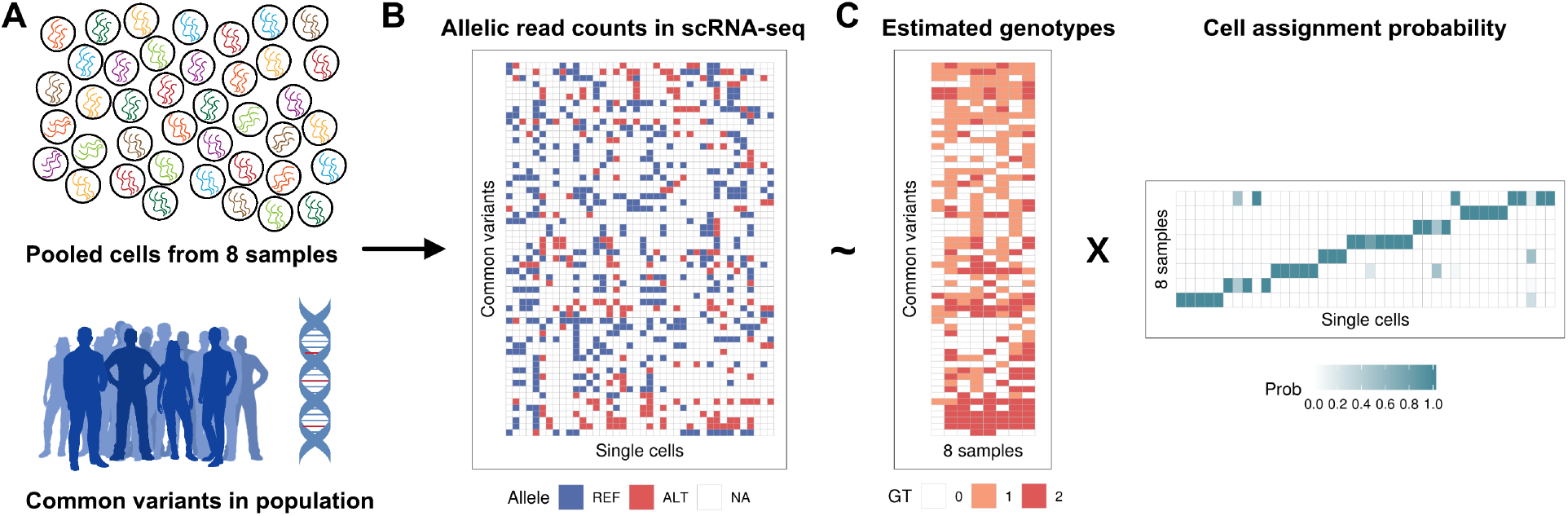
Illustration of Vireo for demultiplexing multi-sample scRNA-seq studies without reference genotype data. (A, B) The inference is based on genotyped common polymorphic variants in each cell, defined based on a standard reference of common human variants. (B, C) The resulting sparse read count matrices of alternative and reference alleles (displayed as compound matrix for simplicity; NA in white denotes no observed reads) are then decomposed into a matrix of estimated genotypes for each input sample and a probabilistic cell assignment matrix.

### Model validation using synthetic data

Initially, we considered synthetic data with a known truth to validate our approach. We considered raw 3’ single-cell RNA-seq data from 10x Genomics platform (v2 kit) for 16 genetically distinct samples from the census of immune cells project that are available from the Human Cell Atlas (Methods) [18]. We then synthetically mixed 8 of these samples (1,000 cells per sample and 4,000 UMIs per cell on average), and simulated 8% of cells as doublets, which were included alongside the sampled singlet cells (“singlets”; Methods). Initially, we evaluated Vireo’s ability to estimate the number of input samples, by comparing the marginal likelihood of multiple Vireo runs assuming increasing numbers of samples in the pool, ranging from six to twelve. Notably, models with at least the true number of input samples (*K* = 8) were evident from an elbow plot of the variational lower bound (Fig. 2A). We also observed that models that assume larger pool sizes (*K* > 8) tended to yield sparse solutions, which means that only the relevant subset of latent samples required to explain the data were used, indicating that the model is robust to reconstructing a larger number of samples than necessary (Supp Fig. S1).

**Figure 2.**
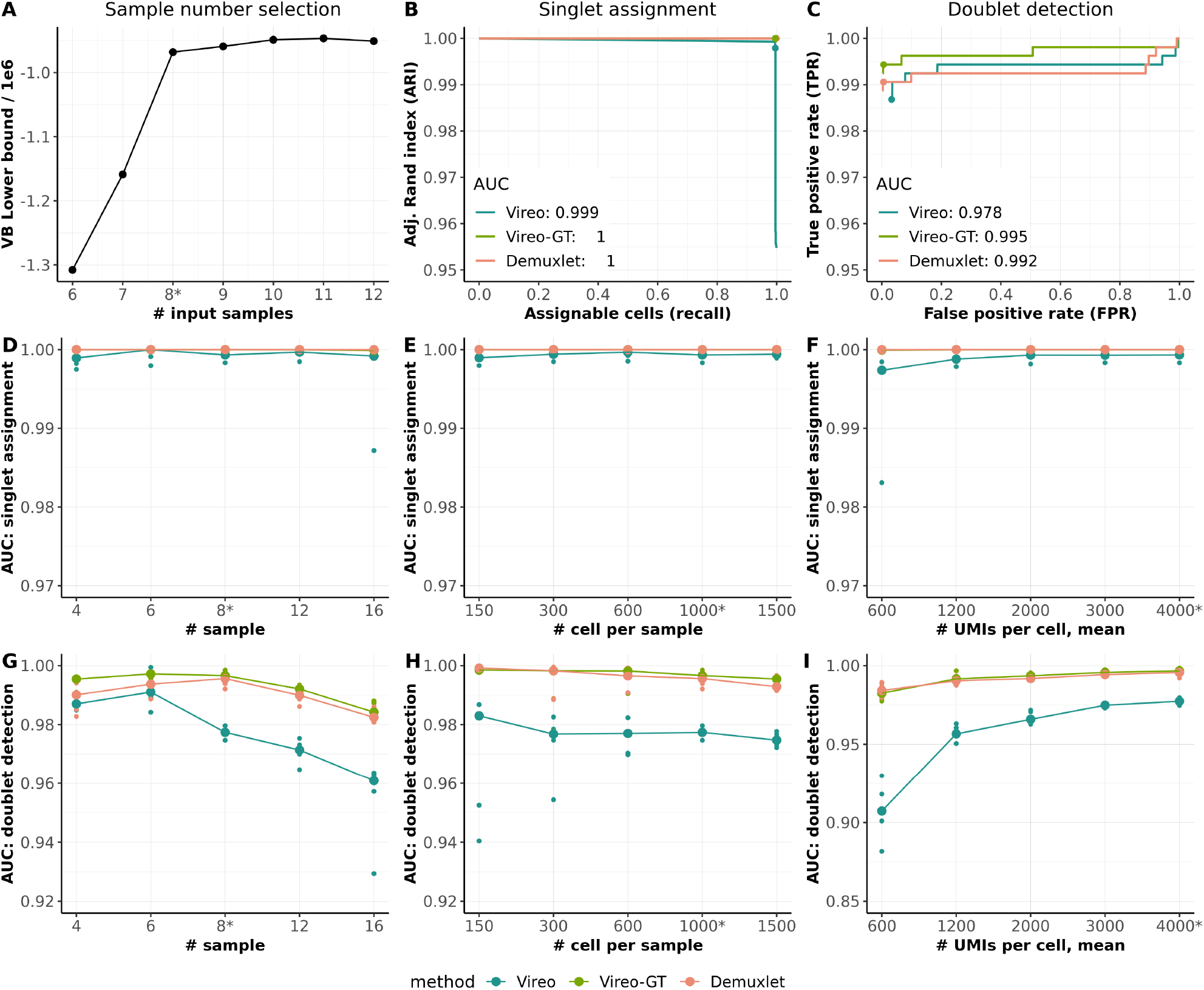
Evaluation of Vireo using synthetic mixtures of up to 16 scRNA-seq datasets, including a comparison to models that require genotype data of the pooled samples (Vireo-GT and Demuxlet). (A-C) Assessment of Vireo performance on one representative simulated dataset consisting of 8 pooled samples with 1,000 cells per sample, 8% doublet rate, and 4,000 UMIs per cell on average. (A) Vireo model evidence (variational lower bound) when varying the pool size assumed in the model. (B) Adjusted Rand index (ARI) between the most likely inferred and the true singlet assignment, when varying the assignment confidence. The recommended cutoff (prob max > 0.9 for Vireo and Vireo-GT, and PRB.SNG1 > 0.67 for Demuxlet) are highlighted as dot. (C) Receiver operating characteristic (ROC) curve for detecting doublets, when varying the assignment confidence. The recommended cutoff (prob doublet > 0.9 for Vireo and Vireo-GT, and 0.67 for Demuxlet) are highlighted with dots. (D-I) Systematic assessment of Vireo and alternative methods on simulated data using a range of parameter choices, using five replicate runs. (D-F) Area under ARI for singlet assignment considering alternative simulation settings, either varying the number of input samples (D), the number of total cells in the dataset (doublet rate varied proportional; 1.2% to 12%; Methods) (E), as well as the number of UMIs per cell (F). (G-I) Area under the ROC for doublet detection, considering the same simulation paramters as in panels D-F. Parameters not varied in either of these experiments were set to their default values (indicated by the star symbol). Small dots in each experiment denote the five replicate simulation experiments, and the big dot denotes the median performance across replicates.

Next, we evaluated the performance Vireo in singlet assignment and doublet detection, where for comparison we also considered alternative models that require full genotype data of the pooled samples (Demuxlet [13] and Vireo-GT, i.e., Vireo with full genotype data; Methods). By measuring the adjusted Rand index (ARI) of the most likely assignment of singlet cells to samples with regard to the true assignments, we found that Vireo achieved markedly accurate results, yielding comparable performance as Vireo-GT and Demuxlet (Fig. 2B). We also varied the assignment confidence (Methods), finding that all three methods achieve near-perfect assignments of the full set of singletons (recall = 1). In the following, we consider the area under the ARI-recall curve (AUC) as a systematic measure for assessing the overall performance of singlet assignment.

Similar to Demuxlet, Vireo can also be used to identify doublet cells, provided that the doublets are formed of combinations of cells from two genetically distinct samples in the pool. Vireo without genotype achieves doublet detection with an overall AUC = 0.978 (e.g, 98.7% sensitivity and 96.7% specificity at prob doublet > 0.9, Fig. 2C), which is only marginally lower than the performance achieved when using genotype data (Vireo-GT or Demuxlet, both AUC ≈ 0.995). In practice, and in the experiments reported below, we recommend prob max >0.9 as the threshold for the singlet assignment, and prob doublet >0.9 for the detection of doublets (see Methods).

Exploring a wider range of settings, we also evaluated the model when varying the number of multiplexed samples (Fig. 2D & 2G), the number of cells sampled in each experiment (Fig. 2E & 2H), and the number of UMIs per cell (Fig. 2F & 2I). As expected, the cell-assignment accuracy decreased with increasing numbers of samples in the pool, but Vireo retained high accuracy for up to 12 multiplexed samples (Fig. 2D & 2G). Beyond 12 samples, there is a risk that the Vireo solution represents a local optima of the optimization of the variational lower bound, which may omit one or multiple samples present in the pool (Fig. 2D). Using current experimental technologies, such high multiplexes are not commonly considered, as high cell counts are associated with greatly increased doublet rates (e.g. on the 10x Chromium platform). Conversely, the accuracy of cell-assignment is consistently high across a larger of cell counts per sample (Fig. 2E), where larger numbers of cells tended to increase accuracy. Similarly, increasing the sequencing coverage resulted in improved accuracy for doublet detection (Fig. 2I), whereas accurate singleton assignments were achieved with extremely low UMIs per cell (Fig. 2F).

Next, we assessed the utility of partial genotype data for a subset of samples in the pool, which as expected increased the model performance, particularly in settings with low sequencing coverage (1,200 UMIs per cell, Supp Fig. S2). We also evaluated the robustness of Vireo when applying the model to biased pools of samples, i.e. settings in which some samples contribute a smaller than expected fraction of cells. Vireo robustly detected and aligned cells to samples with a relative frequency as low as 10% (Supp. Fig. S3), while retaining high accuracy for doublet detection. However, rare sample that were represented by fewer than 100 cells can be missed in some settings.

Finally, we assessed the accuracy of the genotype reconstruction of the pooled samples, finding that Vireo implicitly provides accurate genotype information for expressed variants that are accessible via scRNA-seq, especially for the subset of variants that are covered by at least 10 UMIs per sample (overall precision > 0.96, with heterozygous sites of lowest precision = 0.91; Supp. Fig. S4). Although such estimated genotypic states are intrinsically not available genome-wide, these partial genotype profiles be used as a linking key to align the reconstructed samples to other ’omics data or to combine demultiplexed datasets across experiments (Methods).

### Application to real pooled data

Next, we applied Vireo to two real datasets that have previously been considered to benchmark methods for demultiplexing pooled experiments when genotype information is available for all samples [13].

First, we considered a set of three multiplexed experiments (Fig. 3A-C; W1-W3, between 3,639 and 6,145 cells) of peripheral blood mononuclear cells (PBMCs) from eight lupus patients. We applied Vireo without using genotype information to all cells across these three batches (Methods), thereby also creating an implicit link across all the three experiments. The Vireo cell assignments were markedly consistent with the assignments obtained when using Demuxlet, which, however, depends on genotype data for all samples (Fig. 3A). Similarly, we observed overall concordant doublet cell assignments, although there were larger differences than for singlet assignments (Fig. 3B). We also applied Vireo separately to each of the three datasets and used the inferred genotype state of the samples to link the samples identify across experiments retrospectively (Fig. 3C). This demonstrates how the utility of the inferred genotype data for integrating demultiplexed samples across experiments, which can also be used to link them to other (sequencing-based) assays available from the same samples (Methods).

**Figure 3.**
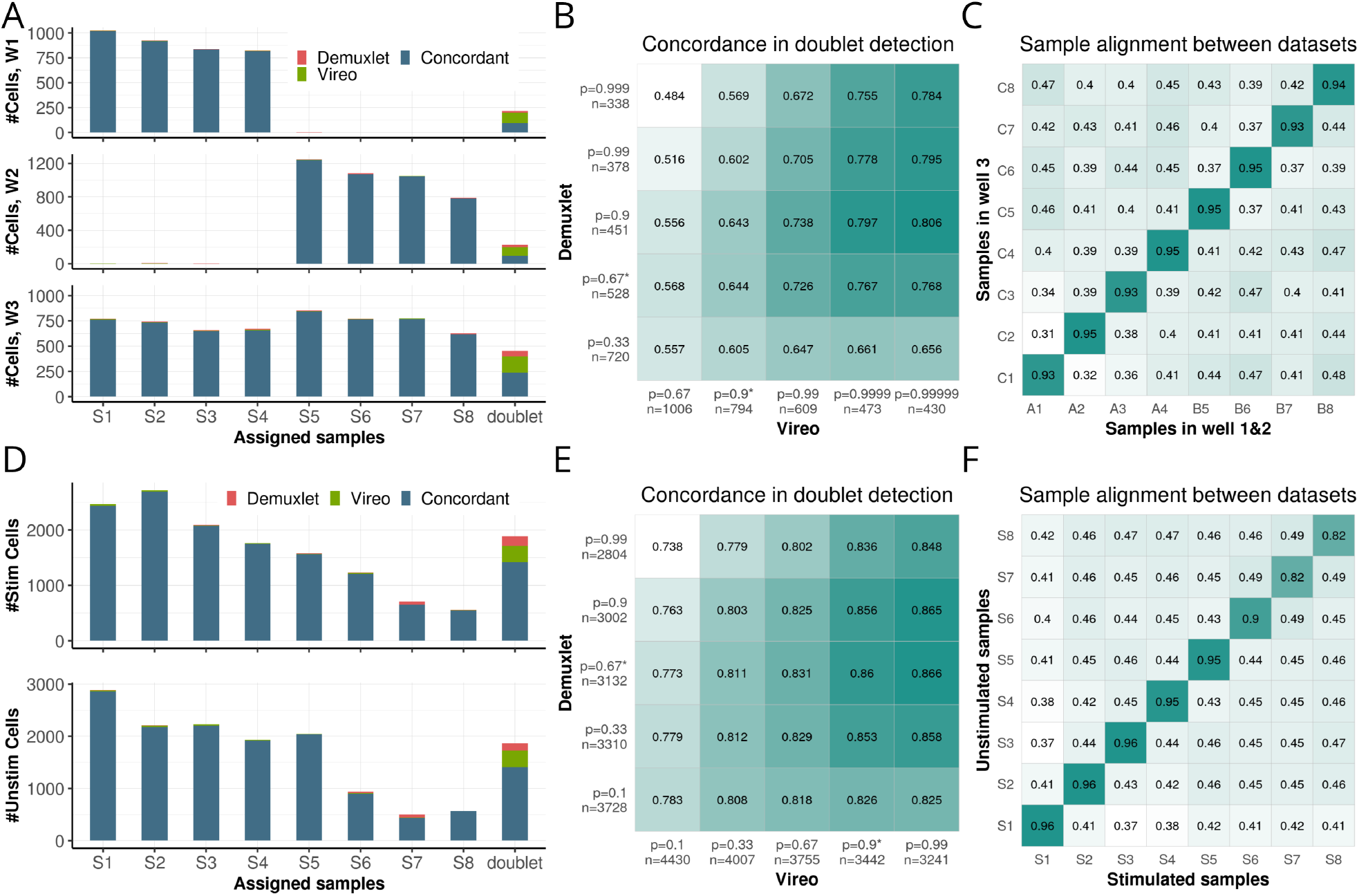
Evaluation of Vireo on data from multiplexed human PBMCs. (A-C) Results obtained on two datasets consisting of two pools of 4 samples each, as well as a third dataset consisting of the union of all 8 samples. (A) Concordance of singlet assignment and doublet detection between Vireo without genotype data and Demuxlet applied with complete genotype reference. Bars denote the number of cells assigned to each sample, either considering cells that were consistently assigned by both Vireo and Demuxlet (blue), or assigned exclusively by Vireo (green) or Demuxlet (red). (B) Concordance of doublet detection between Vireo and Demuxlet when varying the assignment threshold for each method. Note, p denotes the threshold prob doublet in Vireo (x axis) and Demuxlet (y axis) respectively, and n denotes the number of detected doublets. Assignment of cells in A is based on the most probable sample assignment, considering all cells that were not detected as doublet. Cells with a doublet probability (p doublet > 0.9 in Vireo; > 2/3 in Demuxlet) were labelled as doublet cells and are considered in B. (C) Alignment of samples, when applying Vireo separately to the three datasets considered in A. Values in the heatmap denote the fraction of concordant genotype states between pairs of samples from both Vireo runs, considering variants with a read coverage of at least 10 UMIs per sample. (D-F) Results from a second experiment, consisting of two datasets with the same 8 samples pooled in two different conditions: unstimulated and stimulated. Results shown correspond to the panels in (A-C).

As a second use case, we considered two experiments of PBMCs from the same eight patients: one batch with IFN-*β* stimulation and a matched control experiment without stimulus. Cells were cultured for six hours after pooling, which, in contrast to the first dataset, resulted in an imbalanced distribution of cells across samples (Fig. 3D). Despite this distributional bias, Vireo again yielded demultiplexing results that were markedly consistent with the results obtained by methods that require a genotype reference (Fig. 3D-E, and Vireo enabled aligning samples across both experiments (Fig. 3F).

### Leveraging multiplexed designs for differential expression analysis

Finally, we considered the demultiplexed dataset consisting of stimulated and unstimulated cells (Fig. 3D-F) to explore the utility of multi-sample designs for differential gene expression analysis. Graph-based clustering (implemented in Scanpy [19]) applied to the joint dataset consisting of stimulated and unstimulated cells from all eight samples (Fig. 4C) identified eight major clusters, which could be annotated by common cell types (Fig. 4A-B; Supp. Fig. S5). Next, we tested for differential gene expression between the stimulated and unstimulated condition within each cell type (using edgeR, considering cells as replicates [20]). Considering B cells as a representative example (see Supp. Fig. S8-11 for full results), this analysis identified between 78 and 477 DE genes in individual samples (FDR*<*5%; Fig. 4F), with cell count being a major explanatory factor for differences in the number of DE genes (Fig. 4C). Although globally, DE genes tended to be recurrently detected in multiple samples (Fig. 4E), there was a substantial fraction of DE genes that were private to individual samples. For example, the gene OAZ1 (Fig. 4D) was deferentially expressed in four of eight samples, highlighting the importance of inter-individual differences (more examples in Supp. Fig. S6). We also explored carrying out joint testing across all samples (using samples as an explanatory factor in the model in edgeR; Methods), which lead to broadly similar conclusions (Supp. Fig. S7).

**Figure 4.**
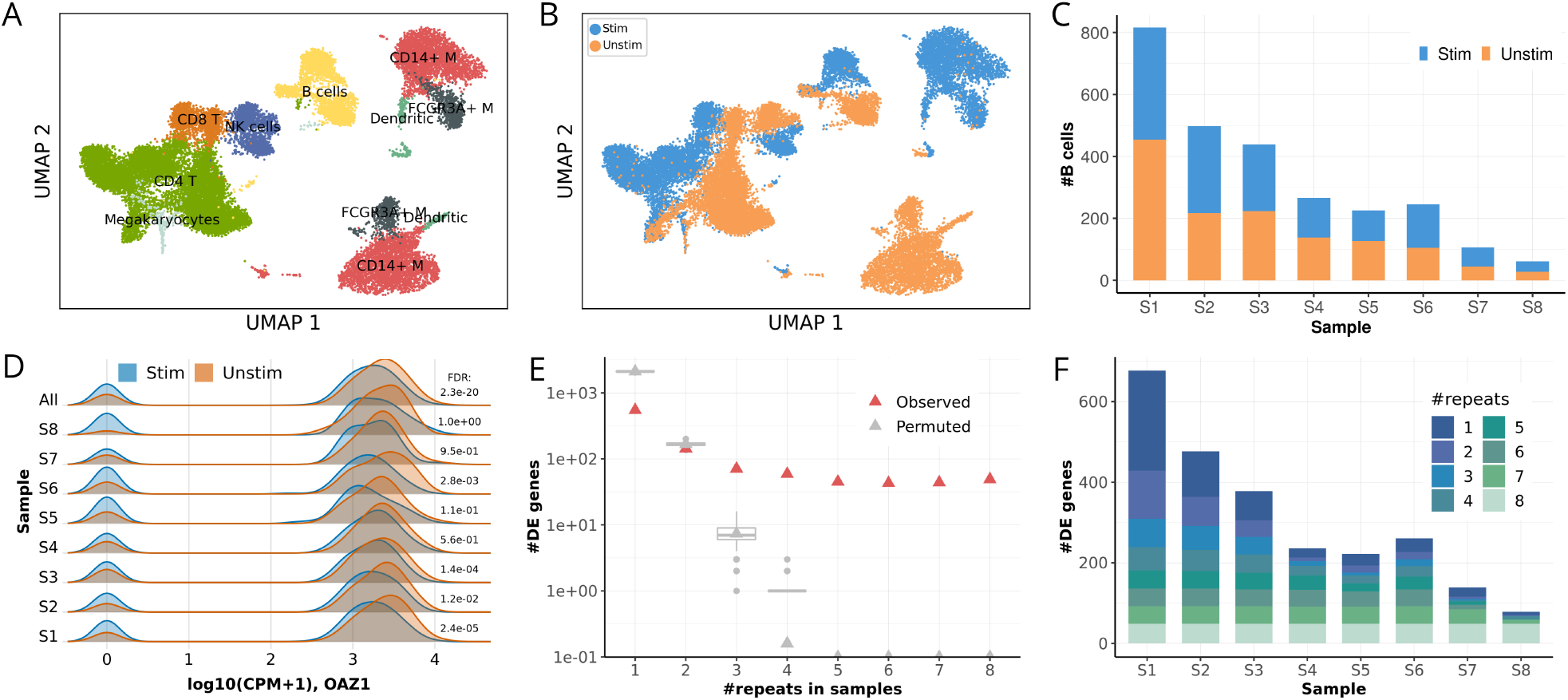
Case study of differential expression analysis of INF-beta stimulated PBMC using two matched pools that consist of 8 samples. (A) UMAP representation of single-cell transcriptome profiles, colored by eight major cell types identified. Note, M denotes Monocytes. (B) Analogous UMAP representation colored by the experimental condition: stimulated versus unstimulated. (C) The number of stimulated and unstimulated B cells sequenced in each of the 8 samples. (D-I) Differential gene expression (DE) analysis between stimulated and unstimulated B cells. (D) The expression level of an example gene OAZ1, depicting the distribution of expression levels in both conditions, either considering each sample separately (S1..S8) or considering aggregated data pooled across all samples (All). FDR: adjusted P value (Benjamini-Hochberg) of each DE test between conditions with likelihood-ratio test. CPM: count per million. (E) Number of recurrently detected DE genes between conditions (FDR< 0.05), detected in at least one to eight samples. Box plots in grey show the recurrence expected by chance (based on 200 permutations). (F) The number of DE genes in each of the 8 samples, categorized by the number of recurrent DE discoveries across samples, that is the number of individuals in which a given gene is identified as DE (FDR< 0.05).

## Conclusion

Here, we have presented Vireo, a Bayesian method for demultiplexing pooled single-cell RNA-seq experiments by exploiting natural genetic barcodes and cell genotyping based on scRNA-seq reads. Uniquely, Vireo does not require any reference genotype data of the specific samples that are pooled in the experiment, while achieving demultiplexing accuracies that are comparable to methods that require a genotype reference. Vireo is implemented using computationally efficient variational Bayesian inference, which provides a fully Bayesian treatment while retaining scalability to large datasets.

Using synthetic mixtures of cells, we have evaluated the accuracy of Vireo for demultiplexing pooled samples, and found it robust to a variety of settings. We also demonstrated the model’s flexibility for handling partial genotype data for some of the samples, should these data be available. Unsurprisingly, we observed that the accuracy of the genotype estimation step per sample is primarily linked to the sequencing coverage, which also substantially affects the ability to detect doublet cells. As the exact requirements for the optimal sequence coverage depend on the cell count and the number of pooled samples, we provide a simulation framework that enables the user to explore parameters thereby aiding the experimental design of pooled studies. If cells from the same individuals are assayed in multiple batches, Vireo can also demultiplex them jointly, which boosts the assignment accuracy, especially in experiments with lower read coverage. Furthermore, the estimated genotypes for individual samples enables aligning samples from the scRNA-seq data with from other ’omics data for the same samples (Fig. 3C), which provides a flexible approach for linking samples across experiments, including multi-omics treatment-control designs.

We noticed that demultiplexing without a genotype reference starts to deteriorate for large sample pools (>12 samples). Increased sequencing coverage may allow for demultiplexing even larger pools, but there remain general experimental limitations for such designs. In particular, as long as the doublet rates scales with increased cell count such designs are not yet of interest.

As future technologies that motivate even larger pool sizes become available, extensions of Vireo that can handle such settings may be warranted. Notably, the demultiplexing accuracy is also linked to read coverage per cell as well as total cell count, two quantities that continues to improve. Thus, the practical limitations on the pool size are likely to increase as single-cell technologies continue to improve.

As a reference-free method, Vireo is particularly useful in settings where samples are treated as biological replicates and the primary object is the variation between samples, which does not require the explicit identification of individual pooled samples(Fig. 3 and 4). Beyond that, Vireo has the intrinsic limitation that the inferred samples cannot be directly identified or linked to metadata. However, when the necessity for sample identity arises, the estimated genotype states are readily available for linking the samples to other ’omics data, e.g., other scRNA-seq batches (Fig. 3C and 3F) or bulk RNA-seq (Supp. Fig. S12). These principles can be applied to any read-based assay, which provides genotypes. Finally, it is straightforward to generate targeted qPCR-based genotyes for a minimal set of discriminatory variants (Supp. Fig. S13). The Vireo software provides helper functions for designing such experiments, which directly leverages the reconstructed genotypes in the pool to define a small set of discrimatory variants (Methods). Molecular barcoding strategies, e.g., [9, 10, 11, 12], have recently emerged as an alternatives to genetic barcoding in many respects courtesy of their more universal applicability. For example, molecular barcoding enables pooling multiple treatment conditions or tissues from the same individual or from individuals with the same genetic background (e.g. inbred model organisms). Nevertheless, the natural genetic barcoding methods, which thanks to Vireo now can be applied even when no genotype data are available, have the advantage of avoiding additional laboratory work, thus reducing the logistical complexity, which can impact processing efficiency and data quality.

## Methods

### Vireo model

Given a list of *N* common variants, we extract allelic expression of these variants in each of *M* cells with RNA-seq data (see below for details on the read pileup approach for variant genotyping). Let *A* and *D* respectively denote the read or UMI count matrices for the alternative allele (i.e., ALT) and the total read depth (i.e., sum of ALT and REF) for *N* variants across *M* cells. Vireo models variation in these counts matrices by employing a clustering model with clusters corresponding to *K* individuals in the pool, with (unknown) genotype states *G*. The values of *G* take on values of 0, 1, or 2, corresponding to homozygous REF, heterozygous and homozygous ALT alleles.

The observed alternative allele counts *A* are modelled as binomial distributed given the read depths

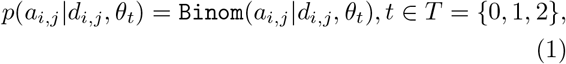

where *t* is the true genotype of variant *i* in cell *j*, and *θ*_*t*_ is the binomial rate parameter that encodes the corresponding allele dosage of the alternative allele for genotype *t*. Theoretically, the allele dosage is *θ*_*t*_ = *t*/2, whereas in practice we allow for deviations to account for sequencing errors, genotype estimation errors, and allelic imbalance.

The genotype in a given cell is defined by a clustering model where the latent genotype *t* for variant *i* in cell *j* is coded by two indicator variables: the cell assignment vector *Z*_*j*_, which assigns cell *j* to a latent sample in the pool, and the genotype identity *G*_*i,k*_, which defines the allelic state of variant *i* in sample *k*. Specifically, the indicator variable *Z*_*j,k*_ = 1 if cell *j* is assigned to sample *k* and 0 otherwise; we also impose the constraint Σ_*k*_ *Z*_*j,k*_ = 1, which means that in expectation each cell originates from exactly one sample. Analogously, the indicator variable *G*_*i,k,t*_ = 1 if the genotype of variant *i* in sample *k* is *t*, and 0 otherwise, and we again require Σ_*t*_ *G*_*i,k,t*_ = 1. The cell assignment matrix *Z* is strictly unknown and needs to be estimated from the observed data. In general, the genotype matrix *G* is also unknown and is estimated jointly with *Z*. If genotype information are available for one or multiple samples in the pool, this information can be encoded as informative prior on *G*; see below.

The likelihood of the full datasets, spanning all *N* variants that were genotyped in each of *M* cells given the cell assignment matrix *Z*, the genotype matrix *G* and binomial parameter ***θ*** follows as:

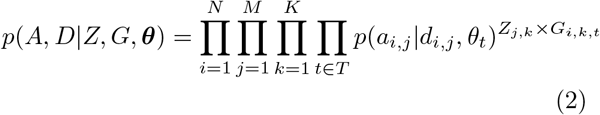

To complete the definition of the model, we introduce prior distributions on the latent variables, which results in the following joint distribution over both observed and latent variables

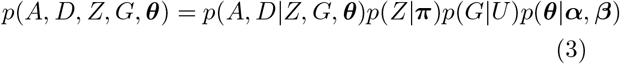

For computational convenience, we use conjugate prior distributions, namely beta distribution for ***θ*** and multinomial distributions for both *Z* and *G*.

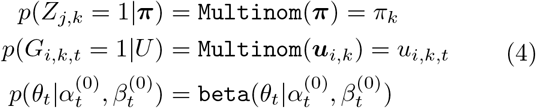

The hyper parameters are constant and set as follows. We use an uninformative prior for *Z*: *π*_*k*_ = 1/*K*, which corresponds to a uniform assignment probability of cells to samples. The user can define other multinomial probabilities, for example to encode known bias in the sample representation. Similarly, we employ a uniform prior on genotype *G*, i.e., *u*_*i,j,t*_ = 1/3 if no genotype data are available. If the genotypes are partially known for a subset of samples and/or variants, a corresponding informative prior is encoded. Specifically, *u*_*i,j,t*_ takes the known genotype value with a relax rate *ξ*, i.e., *u*_*i,j,t*_ = 1 − *ξ* if the known genotype is *t*, otherwise *u*_*i,j,t*_ = *ξ*. The error rate parameter is set to *ξ* = 0.05 by default.

Finally, the hyper parameter for the beta prior on the allelic rate ***θ*** is determined using known germline variants with high coverage: *θ*_0_ ∼ beta(0.3, 29.7), *θ*_1_ ∼ beta(3, 3), and *θ*_2_ ∼ beta(29.7, 0.3), with which the posterior of ***θ*** will be obtained by fitting to the dataset.

### Variational Bayesian inference

Analytical calculation of the posterior distribution of all latent variables given the observed data *p*(*Z, G, **θ**|A, D*) is not tractable. Thus, we consider variational Bayesian inference [21] to obtain an approximate solution, thereby retaining the benefits of a Bayesian treatment while retaining computational scalability to larger scRNA-seq datasets. Briefly, the objective of variational inference is to approximate the exact (intractable) posterior distribution of the latent variables *p*(**Y***|***X**) by a factorized distribution *q*(**Y**) = Π_*i*_ *q*_*i*_(*Y*_*i*_), where **Y** denotes a set of latent variables and **X** denotes the observed variables. The parameters of the variational distribution *q*(**Y**) are determined with the objective to minimise the Kullback-Leibler 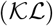 divergence between the approximate distribution *q*(**Y**) and the actual posterior distribution *p*(**Y***|***X**)

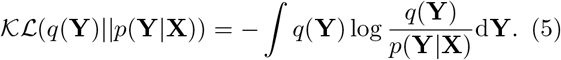

This objective is equivalent to maximizing the lower bound of the full distribution L(*q*), as the log marginal probability of the observed variables is a constant, as follows,

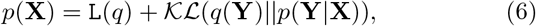

where the lower bound L(*q*) is defined as follows,

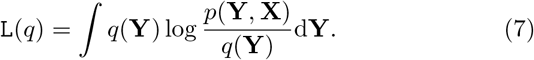

A set of iterative update equations can be derived, which are guaranteed to increase the lower bound

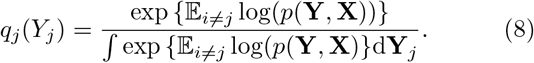

Here, 𝔼_*i≠j*_ denotes an expectation with respect to the distributions *q*_*i*_(**Y**_*i*_) for all *i* ≠ *j*.

For inference in Vireo, we assume a fully factorized distribution *q*(*Z, G, **θ***) = *q*(*Z*)*q*(*G*)*q*(***θ***) to approximate the true posterior distribution *p*(*Z, G, **θ** A, D*), and we assume that *Z* and *G* follow categorical distributions, and ***θ*** follows beta distributions. Based on this assumption, the lower bound can be computed as in Eq(7) (See Supplementary Methods Eq(S1-6)). Following Eq(8), it is possible to derive iterative update equations for the *Q* distribution of the latent variables (see Supplementary Methods Eq (S7-12) for full details).

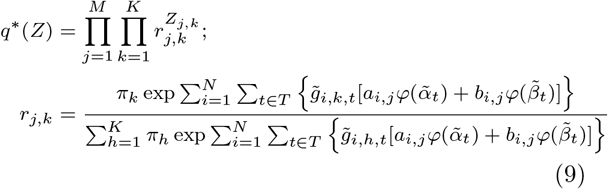

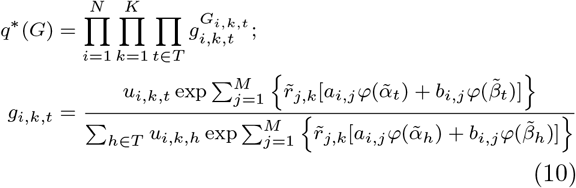

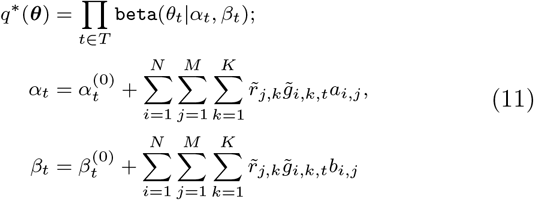

Here, we introduce *b*_*i,j*_ = *d*_*i,j*_ − *a*_*i,j*_ to simplify the notation and *φ*(⋅) denotes the digamma function. To mitigate potential local optima, multiple random restarts are considered (default: 50 restarts) and the solution that maximizes the variational lower bound is selected. Thanks to the implementation with sparse matrix and support of multiple threads, Vireo model is computationally efficient. On a laptop with 16G memory and two 3.5GHz CPUs, Vireo finishes a two-run estimation (see next section) in 6.7 minutes for 14,619 cells in an eight sample pool and 58.1 seconds for 6,145 cells in another eight sample pool (results in Fig. 3).

### Vireo with known genotype or partial genotype

Besides demultiplexing pooled scRNA-seq without any genotype information, Vireo is also able to leverage any available genotype information. In the case that the genotype is available for all pooled samples, we only use the variants with known genotype and set the genotype probability variable *G* as known and fixed, which can be derived from the GT tag (for categorical genotype), GP or GL tag (genotype probability or likelihood) in the VCF file. By default, we use GT as it is the most commonly available tag.

Alternatively, Vireo also supports the use of any partial genotypes via a two-step run approach. In the first run, Vireo does not use any genotype information but infers the genotype for each sample. Then, we align the samples with known genotype to these identified samples in this run and replace the estimated genotype probability with the input known values. Therefore, we obtain a genotype probability matrix with mixed known and inferred samples, which we then use as a prior of *G*, instead of the default uniform prior in the second run. Finally, we report the results of the second run as the result of Vireo.

### Estimation of the number of pooled samples

Access to the variational lower bound (Eq. 7) allows for estimating the number of samples in a given pool. Briefly, by comparing alternative Vireo runs with increasing numbers of samples it is possible to identify the most probable value with the elbow plot (e.g., Fig. 2A), which provides an objective means to define this parameter.

A second strategy is to set a large number of samples and prune some of the samples posthoc, as variational Bayes model is self-regularizing and hence avoids over-fitting (see Supp. Fig. S1). In practice this approach can also increase robustness as the effective number of samples in a pool can be larger than anticipated due to doublets; see Section below.

### Multiple random initializations

When genotype is not given, Vireo uses a pre-step with multiple random initializations to avoid local optima. By default, Vireo runs for 50 random initializations, each with a short iterations (15 by default). Then the initialization with highest log likelihood will be continued.

As discussed in the above subsection, another strategy to find all *K* pooled samples is searching from a larger number of clusters. By default, we search 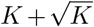 clusters in this pre-step, and only keep the *K* clusters with largest number of assigned cells to continue, and discarded the 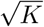 smaller clusters.

### Doublet detection

To detect doublets, we construct the genotype of each pair of samples and expand the *K* biological samples by introducing in additional (*K* − 1) * *K*/2 doublet competitors. For simplicity, we assume that the genotype of a doublet sample can be described as the average between two combined samples. Specifically, for a given variant and genotype probability vectors for two samples are ***x*** = [*x*_0_, *x*_1_, *x*_2_] and ***y*** = [*y*_0_, *y*_1_, *y*_2_], we define the expected genotype for the doublet sample as follows,

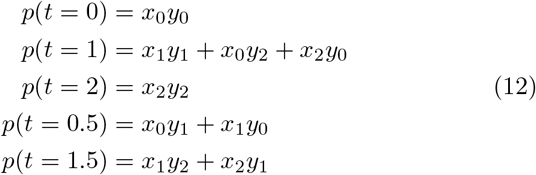

where we introduce two pseudo-genotypes *t* = 0.5 and *t* = 1.5 respectively for combinations of genotype 0 & 1 and 1 & 2 in the doublet sample. For convenience, we consider the binomial parameters for the alternative allelic reads and assume that the binomial parameters *θ*_0.5_ and *θ*_1.5_ also follow beta distributions. We approximate the hyper-parameters of the beta distribution empirically by respectively taking the ratio and shapes with the arithmetic and geometric means from the two ordinary genotypes. The resulting distribution of *θ*_0.5_ can be expressed as follows

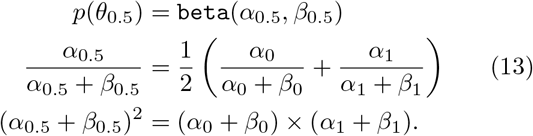

Similarly, we define the distribution of *θ*_1.5_.

In this augmented model, we have the full distribution for the extended genotype reference *G* and ***θ***, consisting of *K* biological and (*K* − 1) * *K*/2 doublet samples. In this model we can calculate the probability that a cell originates from one of the doublet samples using Eq(8).

As an additional refinement, we specify a nonuniform prior on *η* to define the *a priori* believe of observing a doublet. Specifically, the prior binomial distribution ***π*** is constructed as follows

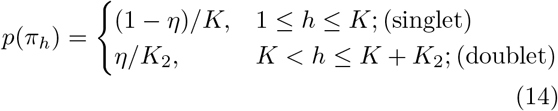

where *K*_2_ = (*K* − 1) × *K*/2 as number of the combined sample pairs. The prior probability for doublet cells is low in most assays, e.g., *η* = 0.05. In case of the 10X chromium platform, the prior value can be estimated as a function of the number of loaded cells *M*, e.g. *η* = *M*/100000 following [13], which by default are used in our experiments.

Therefore, we can obtain the posterior of each cell’s sample identity, i.e., the probability of cell *j* coming from any of the *K* input samples or *K*_2_ combined sample pairs (i.e., doublet). We use the highest assignment probability of the *K* input samples, prob max, as the confidence score for singlet assignment and use the summarised probability of all *K*_2_ sample pairs as the confidence score of a doublet, namely prob doublet.

### Alignment of samples between multiple data sets

Vireo implicitly estimates the genotypes for subset of variants with sufficient coverage (good accuracy for variants with >10 reads per sample; see Supp. Fig. S4). Among other use cases, these estimated genotypes allow for aligning scRNA-seq profiles from samples in a pool to other ’omics data by matching genotype profiles.

When Vireo is applied to multiple data sets that consist of the same samples, the estimated genotype also allows for aligning samples across data sets (e.g. Fic. 3C and 3F for multiple pools, and Supp. Fig. S12 for multiple ’omics). The software implementation of Vireo provides support functions for this step by calculating the fraction of variants with matched genotype between two or multiple experiments.

### Identification of discriminatory variants

Given a set of variants for which estimated genotypes are available, the Vireo software implements a heuristic to define a minimal and informative set of discriminatory variants. This set of variants can be used to perform qPCR-based genotyping or for other targeted genoytping methods. Briefly, the algorithm implemented in Vireo prioritises variants with largest information gain in splitting samples, as follows.

1. Remove variants with *<* 20 UMIs per sample.
2. Initialize the variant set *S* = {}, and the split *T* among *K* samples, and calculate the initial entropy *H*(*T*) = 0
3. Rank variants by the information gain *IG*(*T, v*) = *H*(*T*) − *H*(*T|v*)
4. Select the variant with highest information gain and update *S*, *T*, and *H*(*T*)
5. If *H*(*T*) = log_2_(*K*), return *S* and *T*, otherwise go to step 3.

Additionally, variants with homozygous alternative allele in the pooled samples can also be filtered out before hand if needed. Examples of discriminatory variant sets for the six-sample pool from HipSci project are shown in Supp. Fig. S13.

### Differential expression analysis

Differential expression analysis was performed with edgeR [20] between stimulated and control samples (Fig. 4 and Supp. Fig. S7-11). A generalised regression is applied in edgeR to test whether the stimulation contributes to the expression variation on a certain gene by using a likelihood-ratio test. By using the raw UMI count, we performed cell type specific DE analysis with the following three different strategies for all cells jointly (Supp. Fig. S7).

Method 1: *y* ∼ ~ cdr + condition + sample, where *y* is the expression count for a specific gene, which is regressed three covariants: cdr, the cell detection rate (i.e., the fraction of expressed gene in each cell), stimulation condition and the sample identity.

Method 2 and 3: *y* cdr+condition, where we ignore the sample identity of each cell in the pool (Method 2). This same model can also be used in a pseudo-bulk manner with summarize the count for all cells in the same type in a sample (Method 3). Alternatively, we can always perform this model at single cell level for each sample separately (Fig. 4D-F).

### ScRNA-seq data from Demuxlet paper

In this study, we considered two existing multiplexed scRNA-seq datasets that consist of a total of five batches [13]. Raw .bam files were obtained from the Gene Expression Omnibus (GEO; accession number GSE96583). The processed results from Demuxlet for these five batches were directly downloaded from https://github.com/yelabucsf/demuxlet_paper_code. Approximately 37 million common variants (allele frequency > 0.0005) extracted from the 1000 Genome Project, phase 3 [17] were used as candidate variants for scRNA-seq genotyping. We provide an companion Python package `cellSNP` (https://pypi.org/project/cellSNP) for this task, which enables generating selected pile-ups from scRNA-seq data. We discarded non-bi-allelic variants as well as variants with fewer than 20 total UMIs across all cells or minor (i.e., second) allele has less than 10% of total UMIs. The final output of `cellSNP` are two variantsby-cells matrices, *A* and *D*, for UMI counts of alternative allele and the total counts respectively, which are used as input for the Vireo model.

### Bulk RNA-seq and scRNA-seq from HipSci project

In order to link the inferred samples to other ’omics data, we used one scRNA-seq pool for iPSC differentiation in HipSci project (10x Genomics platform, experiment 44, day 0) with six samples: *pipw*, *jejf*, *qehq*, *juuy*, *uilk*, and *toco* [15], and their according bulk RNA-seq data for each sample [22] (http://www.hipsci.org). Both scRNA-seq and bulk RNA-seq data sets were downloaded in .bam files and genotyped on 7.4 millions common bi-allelic variants (minor allele frequency >5%) extracted from the 1000 Genome Project with `cellSNP` package. For single-cell data, we only keep variants with minor allele frequency ≥ 0.1 and ≥ 20 UMIs. For each bulk RNA-seq sample, we also only keep variants with minor allele frequency ≥ 0.1 but require ≥ 100 read counts. Then the genotypes of each bulk RNA-seq sample can be used to align to the samples that are demultiplexed from scRNA-seq data.

### Synthetic data

We obtained raw 3’ scRNA-seq data based on 10x Genomics platform (v2 kit) for 16 genetically distinct samples from Human Cell Atlas (Census of Immune Cells) [18]. These data set are not pooled and each sample has its own sequencing run. We only used data from the first channel (each sample with around 100 millions reads), which is in the range of a standard 10x sequencing run. We first mapped the raw fastq files to the human genome hg38 by CellRanger v2.1 provided by 10x Genomics (cellranger count command line). Then we used `cellSNP` to genotype 7.4 millions common variants (minor allele frequency >5%) extracted from the 1000 Genome Project for these 16 samples in a pseudo-bulk manner. We only keeps variants with 1) >100 UMIs summarised across 16 samples, 2) >10% UMIs from minor allele, and 3) *<*5 UMIs for other alleles (i.e., not annotated reference and alternative alleles). Therefore, we obtained the genotypes of 62,193 variants for these 16 samples, which are feed into Demuxlet and Vireo-GT.

By only keeping cells with >500 genes and >1000 UMIs, we had in total 66,410 cells across 16 samples, with each sample having 2,495 to 4,909 cells. On average, there are 4,000 UMIs per cell (median 2,700 UMIs). In the synthetic mixture, we pooled reads for a subset of cells from each sample (in bam format, aligned reads) and generated multiplexed scRNA-seq data (also in bam format). The script to generating these synthetic data is provided in Vireo’s GitHub repository. Doublets were added into the pooled data by adding proportional extra cells and combine them with another cells randomly. The doublet rate is *N/*100, 000 where *N* is the total number of cells in the pool.

By default, we pooled 1000 cells from each of 8 samples with doublet rate of 8%. This simulator also allows setting different size of input samples, for example by setting one sample with fewer cells ranging from 50 to 500 (Supp. Fig. S3). With the synthetic data in bam format, we can even further sub sample reads by using `samtools` with -s argument, e.g., 15-75% in Fig 2F.

All these simulations were randomly repeated for five times to account for the variability in the simulation.

## ABBREVIATIONS

scRNA-seq: single-cell RNA-seq
SNP: Single-nucleotide polymorphism
AUC: area under the curve
ARI: adjusted Rand index

## Availability of Data and Materials

Vireo model has been implemented as a standard Python package, which is freely available at https://pypi.org/project/vireoSNP with Apache License 2.0. All scripts to replicate the simulations in this paper are also included in linked GitHub repository. Vireo’s manual with examples is available at https://vireoSNP.readthedocs.io.

## Competing interests

The authors declare that they have no competing interests.

## Authors’ contributions

O.S. and D.J.M conceived and guided the study. Y.H. developed and implemented the model. Y.H., D.J.M and O.S. carried out the experiments. Y.H., O.S. and D.J.M interpreted the results and wrote the paper.

## Acknowledgements

We would like to thank Raghd Rostom, Maria Ban, Marc Jan Bonder, Stephen Sawcer and Sarah Teichmann for fruitful discussions.

## Ethics approval and consent to participate

Not applicable.

## Additional File

Additional file 1 — Supplementary Methods and Supplementary Figures S1–S12.

