## Supplementary materials for "Vireo: Bayesian demultiplexing of pooled single-cell RNA-seq data without genotype reference"

[1]Department of Clinical Neurosciences, University of Cambridge, CB2 0QQ, Cambridge, UK; [2]EMBL-European Bioinformatics Institute, Wellcome Genome Campus, CB10 1SD, Hinxton, Cambridge, UK; [3]St Vincent's Institute of Medical Research, Fitzroy, Victoria 3065, Australia; [4]Melbourne Integrative Genomics, University of Melbourne, Parkville, Victoria 3010, Australia; [5]European Molecular Biology Laboratory, Genome Biology Unit, 69117 Heidelberg, Germany; [6]Division of Computational Genomics and Systems Genetics, German Cancer Research Center (DKFZ), 69120, Heidelberg, Germany.

[*]Corresponding authors.

### 1 Supplementary methods

In this supplementary section, we will first re-introduce the notation we use, then derive the detailed computation of the lower bound of the variational distribution $\mathtt{L}(q)$ in Eq(7) in the main text, and lastly derive the updates of each variational component in equations Eq(9-11) in the main text. By leveraging the read counts of alternative alleles $A$ and both alleles (namely depth) $D$ from $N$ variants in $M$ cells, Vireo aims to estimate the joint posterior distribution of sample identity $Z$ for each cell $j$ from each sample $k$, the genotype $G$ for variant $i$ in each sample $k$, and the corresponding alternative allele rate $\boldsymbol{\theta}$ for each genotype $t \in \{0, 1, 2\}$. As described in the main text, we used multinomial priors for the categorical variables $Z$ and $G$ with hyper-parameters $\boldsymbol{\pi}$ and $U$, respectively, and by default both take uniform multinomial priors. We used beta priors for the parameter of the alternative allele rate $\boldsymbol{\theta}$, and we took the hyper-parameters $(\alpha_t, \beta_t), t \in \{0, 1, 2\}$ that generally fit well to highly expressed germline variants in standard scRNA-seq data set (not multiplexed). Specifically, the default prior distribution are: $\theta_0 \sim \mathtt{beta}(0.3, 29.7)$, $\theta_1 \sim \mathtt{beta}(3, 3)$, and $\theta_2 \sim \mathtt{beta}(29.7, 0.3)$.

Next, the lower bound $\mathtt{L}(q)$ in Eq(7) can be written as follows

$$\begin{aligned}\mathcal{L}(q) =& \sum_Z \sum_G \int_{\boldsymbol{\theta}} q(Z, G, \boldsymbol{\theta}) \log \left\{ \frac{p(A, D, Z, G, \boldsymbol{\theta})}{q(Z, G, \boldsymbol{\theta})} \right\} \mathrm{d}Z \mathrm{d}G \mathrm{d}\boldsymbol{\theta} \\ =& \mathbb{E}_{G,Z,\boldsymbol{\theta}}[\log p(A, D, Z, G, \boldsymbol{\theta})] - \mathbb{E}_{G,Z,\boldsymbol{\theta}}[\log q(Z, G, \boldsymbol{\theta})] \\ =& \mathbb{E}_{G,Z,\boldsymbol{\theta}}[\log p(A, D|Z, G, \boldsymbol{\theta})] + \mathbb{E}_Z[\log p(Z|\boldsymbol{\pi})] + \mathbb{E}_G[\log p(G|U)] + \\ & \mathbb{E}_{\boldsymbol{\theta}}[\log p(\boldsymbol{\theta}|\boldsymbol{\alpha}, \boldsymbol{\beta})] - \mathbb{E}_Z[\log q(Z)] + \mathbb{E}_G[\log q(G)] + \mathbb{E}_{\boldsymbol{\theta}}[\log q(\boldsymbol{\theta})]\end{aligned} \tag{S1}$$

where each part is expressed below.

$$\mathbb{E}_{G,Z,\boldsymbol{\theta}}[\log p(A, D|Z, G, \boldsymbol{\theta})] = \sum_{i=1}^{N} \sum_{j=1}^{M} \sum_{k=1}^{K} \sum_{t \in T} \left\{ \tilde{r}_{j,k} \tilde{g}_{i,k,t} \binom{d_{i,j}}{a_{i,j}} [a_{i,j} \varphi(\tilde{\alpha}_t) + b_{i,j} \varphi(\tilde{\beta}_t)] \right\} \tag{S2}$$

$$\mathbb{E}_{\boldsymbol{\theta}}[\log p(\boldsymbol{\theta}|\boldsymbol{\alpha}, \boldsymbol{\beta})] = \sum_{t \in T} (\alpha_t + \beta_t - 2)\varphi(\tilde{\alpha}_t + \tilde{\beta}_t) - (\alpha_t - 1)\varphi(\tilde{\alpha}_t) - (\beta_t - 1)\varphi(\tilde{\beta}_t) - \log(\mathrm{B}(\alpha_t, \beta_t)) \tag{S3}$$

$$\mathbb{E}_{\boldsymbol{\theta}}[\log q(\boldsymbol{\theta}|\tilde{\boldsymbol{\alpha}}, \tilde{\boldsymbol{\beta}})] = \sum_{t \in T} (\tilde{\alpha}_t + \tilde{\beta}_t - 2)\varphi(\tilde{\alpha}_t + \tilde{\beta}_t) - (\tilde{\alpha}_t - 1)\varphi(\tilde{\alpha}_t) - (\tilde{\beta}_t - 1)\varphi(\tilde{\beta}_t) - \log(\mathrm{B}(\tilde{\alpha}_t, \tilde{\beta}_t)) \tag{S4}$$

$$\mathbb{E}_Z[\log p(Z|\boldsymbol{\pi})] = \sum_{j=1}^{M}\sum_{k=1}^{K}\{\tilde{r}_{j,k}\log(\pi_k)\}, \quad \mathbb{E}_G[\log p(G|U)] = \sum_{i=1}^{N}\sum_{k=1}^{K}\sum_{t\in T}\{\tilde{g}_{i,k,t}\log(u_{i,t})\} \tag{S5}$$

$$\mathbb{E}_Z[\log q(Z)] = \sum_{j=1}^{M}\sum_{k=1}^{K}\{\tilde{r}_{j,k}\log(\tilde{r}_{j,k})\}, \quad \mathbb{E}_G[\log q(G)] = \sum_{i=1}^{N}\sum_{k=1}^{K}\sum_{t\in T}\{\tilde{g}_{i,k,t}\log(\tilde{g}_{i,k,t})\} \tag{S6}$$

Note, the variables with tilde hat are the estimated parameters otherwise are fixed hyper parameters, including $\alpha_t$ and $\beta_t$. Same below.

Then, following the general updating rule in the mean-field variational inference (see main text Eq(8)), we can update the parameters in each component alternately while fixing all other components of the variational distributions and reach the finalized equations Eq(9-11) in the main paper.

First, by using the distributions of genotype $G$ and alternative allele rate $\boldsymbol{\theta}$ that are estimated from a previous step in the iteration, we can analytically update the distribution of the sample assignment $Z$ by a categorical distribution.

$$\begin{aligned}\log q^*(Z) &= \mathbb{E}_{G,\boldsymbol{\theta}}[\log p(A,D,Z,G,\boldsymbol{\theta})] + \texttt{const.} \\ &= \sum_{j=1}^{M}\sum_{k=1}^{K}\sum_{i=1}^{N}\sum_{t\in T} Z_{j,k}\left\{\tilde{g}_{i,k,t}[a_{i,j}\varphi(\tilde{\alpha}_t) + b_{i,j}\varphi(\tilde{\beta}_t)]\right\} + \texttt{const.}\end{aligned} \tag{S7}$$

where $\varphi(\cdot)$ is the digamma function, the same below. As $q(Z_j)$ for any $j$ follows a multinomial distribution, we can therefore have the updated parameter $r_{j,k}$, namely the probability of cell $j$ from component $k$ as follows,

$$r_{j,k} = \frac{\pi_k \exp\sum_{i=1}^{N}\sum_{t\in T}\left\{\tilde{g}_{i,k,t}[a_{i,j}\varphi(\tilde{\alpha}_t) + b_{i,j}\varphi(\tilde{\beta}_t)]\right\}}{\sum_{h=1}^{K}\pi_h \exp\sum_{i=1}^{N}\sum_{t\in T}\left\{\tilde{g}_{i,h,t}[a_{i,j}\varphi(\tilde{\alpha}_t) + b_{i,j}\varphi(\tilde{\beta}_t)]\right\}} \tag{S8}$$

Second, with a similar procedure, the analytical updates for the genotype distribution can be written in the form of a categorical distribution as follows,

$$\begin{aligned}\log q^*(G) &= \mathbb{E}_{Z,\boldsymbol{\theta}}[\log p(A,D,Z,G,\boldsymbol{\theta})] + \texttt{const.} \\ &= \sum_{i=1}^{N}\sum_{k=1}^{K}\sum_{t\in T}\sum_{j=1}^{M} G_{i,k,t}\left\{\tilde{r}_{j,k}[a_{i,j}\varphi(\tilde{\alpha}_t) + b_{i,j}\varphi(\tilde{\beta}_t)]\right\} + \texttt{const.}\end{aligned} \tag{S9}$$

where the updated probability of variant $i$ in component $k$ equals to $t$ can be expressed as follows,

$$g_{i,k,t} = \frac{u_{i,k,t}\exp\sum_{j=1}^{M}\left\{\tilde{r}_{j,k}[a_{i,j}\varphi(\tilde{\alpha}_t) + b_{i,j}\varphi(\tilde{\beta}_t)]\right\}}{\sum_{h\in T} u_{i,k,h}\exp\sum_{j=1}^{M}\left\{\tilde{r}_{j,k}[a_{i,j}\varphi(\tilde{\alpha}_h) + b_{i,j}\varphi(\tilde{\beta}_h)]\right\}}. \tag{S10}$$

Lastly, the analytical updates of the distribution of the alternative allele rate $\boldsymbol{\theta}$ can be expressed in the form of a beta distribution as follows,

$$\begin{aligned}\log q^*(\boldsymbol{\theta}) &= \mathbb{E}_{G,Z}[\log p(A,D,Z,G,\boldsymbol{\theta})] + \texttt{const.} \\ &= \sum_{t\in T}\sum_{i=1}^{N}\sum_{j=1}^{M}\sum_{k=1}^{K}\{\tilde{r}_{j,k}\tilde{g}_{i,k,t}[a_{i,j}\log(\theta_t) + (d_{i,j} - a_{i,j})\log(1-\theta_t)]\} + \texttt{const.} \\ &= \log(\texttt{beta}(\theta_t|\alpha_t,\beta_t))\end{aligned} \tag{S11}$$

where the parameters for this beta distribution are

$$\alpha_t = \sum_{i=1}^{N}\sum_{j=1}^{M}\sum_{k=1}^{K}\left\{\tilde{r}_{j,k}\tilde{g}_{i,k,t}a_{i,j}\right\};\beta_t = \sum_{i=1}^{N}\sum_{j=1}^{M}\sum_{k=1}^{K}\left\{\tilde{r}_{j,k}\tilde{g}_{i,k,t}d_{i,j}\right\}. \tag{S12}$$

Now, by updating these parameters iteratively, we can achieve the maximized lower bound of $\mathtt{L}(q)$, and equivalently the minimized KL divergence $\mathrm{KL}(q(Z,G,\boldsymbol{\theta})||p(Z,G,\boldsymbol{\theta}|A,D))$.

### 2 Supplementary Figures

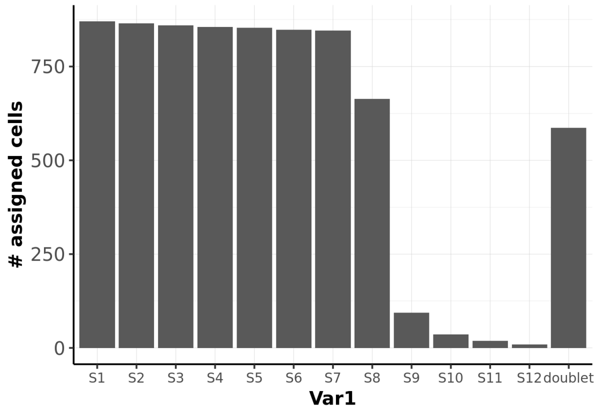


Figure S1: The distribution of number of cells assigned to individual samples when running Vireo assuming a too large pool of size N=12. The true number of samples in the pool is N=8.

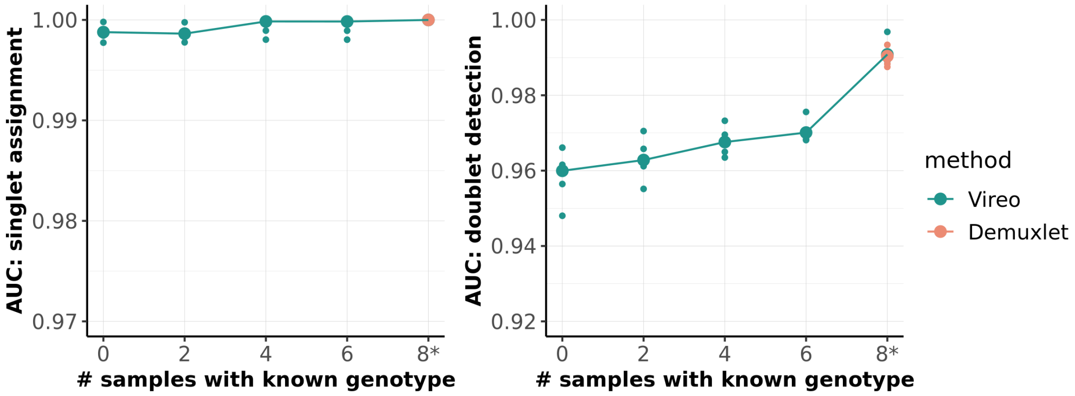


Figure S2: AUC for doublet detection (ROC curve) and singlet assignment (ARI-recall curve) when varying the number of samples with known genotype. There are 8 samples in pool, with 1000 cells per sample and 1,200 UMIs per cell, instead of the default 4,000 UMIs. The five small dots in each category denote the five simulation replicates, and the big dot denotes the median value of the five.

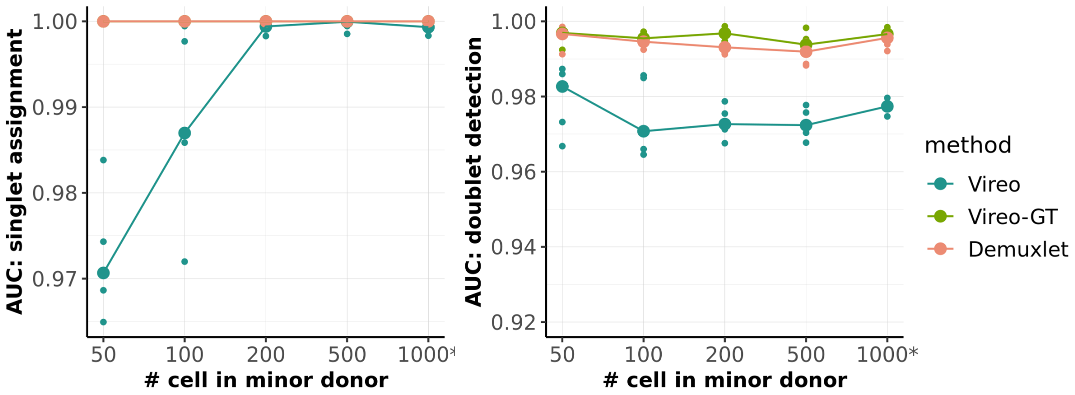


Figure S3: AUC for doublet detection (ROC curve) and singlet assignment (ARI-recall curve) when varying the cell number of the one minor sample in the pool. There are 8 samples in the pool, and the other 7 samples have 1000 cells each. The five small dots in each category denote the five simulation replicates, and the big dot denotes the median value of the five.

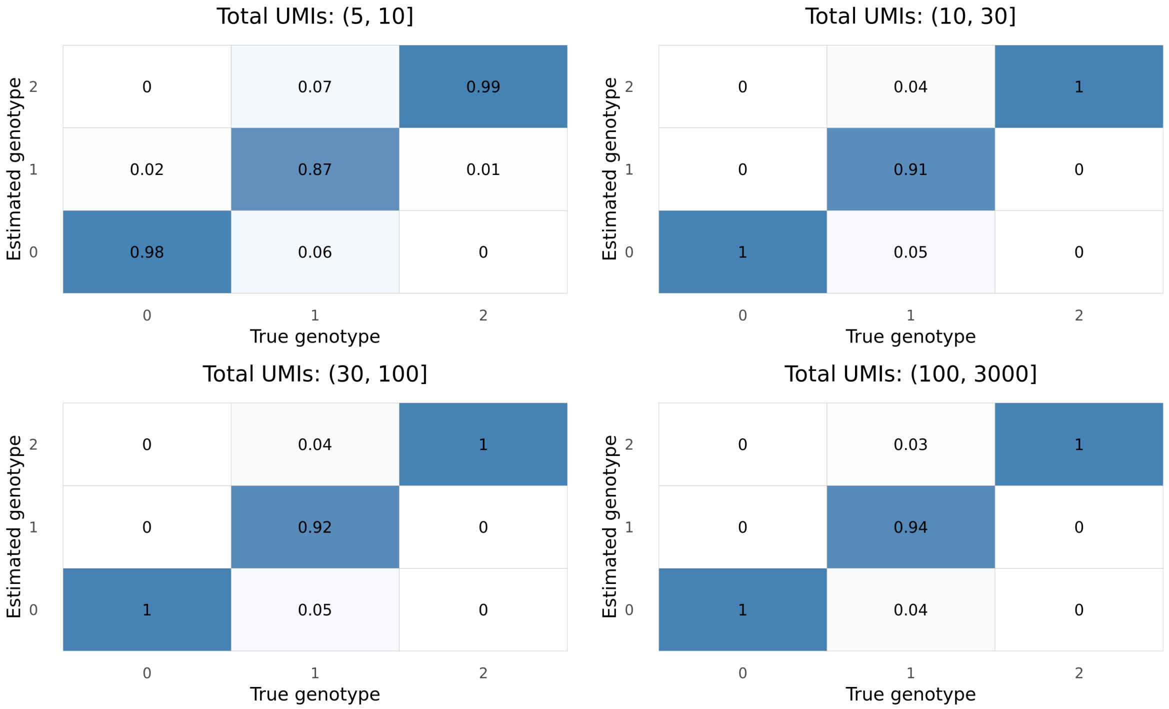


Figure S4: Confusion matrix of genotype estimate for variants with different total UMIs per sample. The estimated genotype is the one with highest probability.

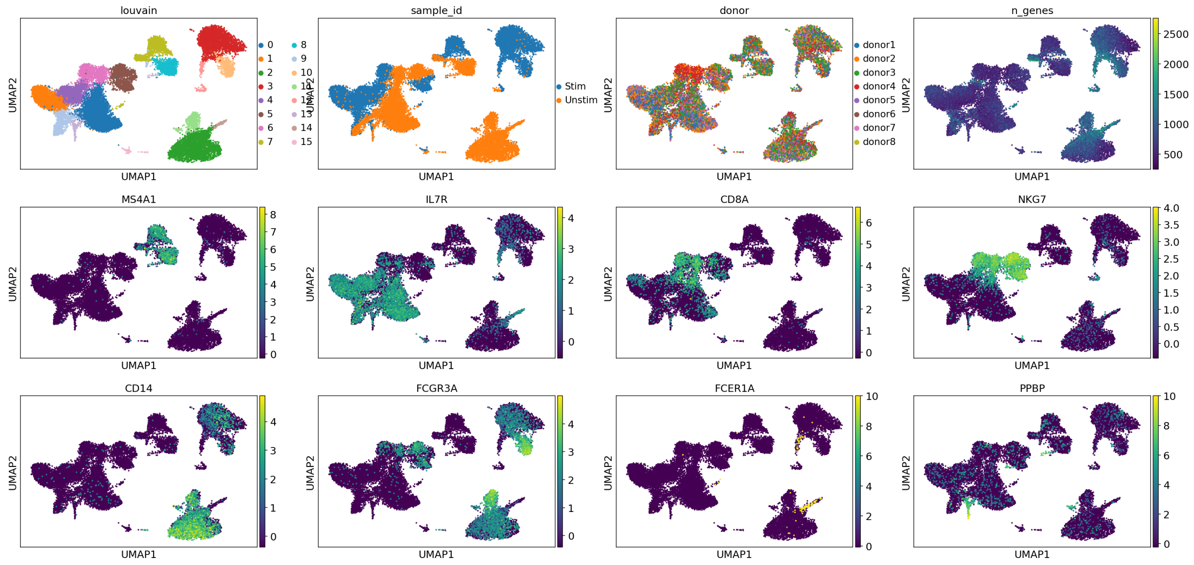


Figure S5: Cell type identification with clustering transcriptome and visualised with UMAP plot. Initially, the stimulated and unstimulated cells were clustered into 16 groups, considering the condition variation. Then, the 16 groups were manually merged into eight cell types by exploring the cell-type marker genes. The final eight cell types are presented in the main Fig 4A.

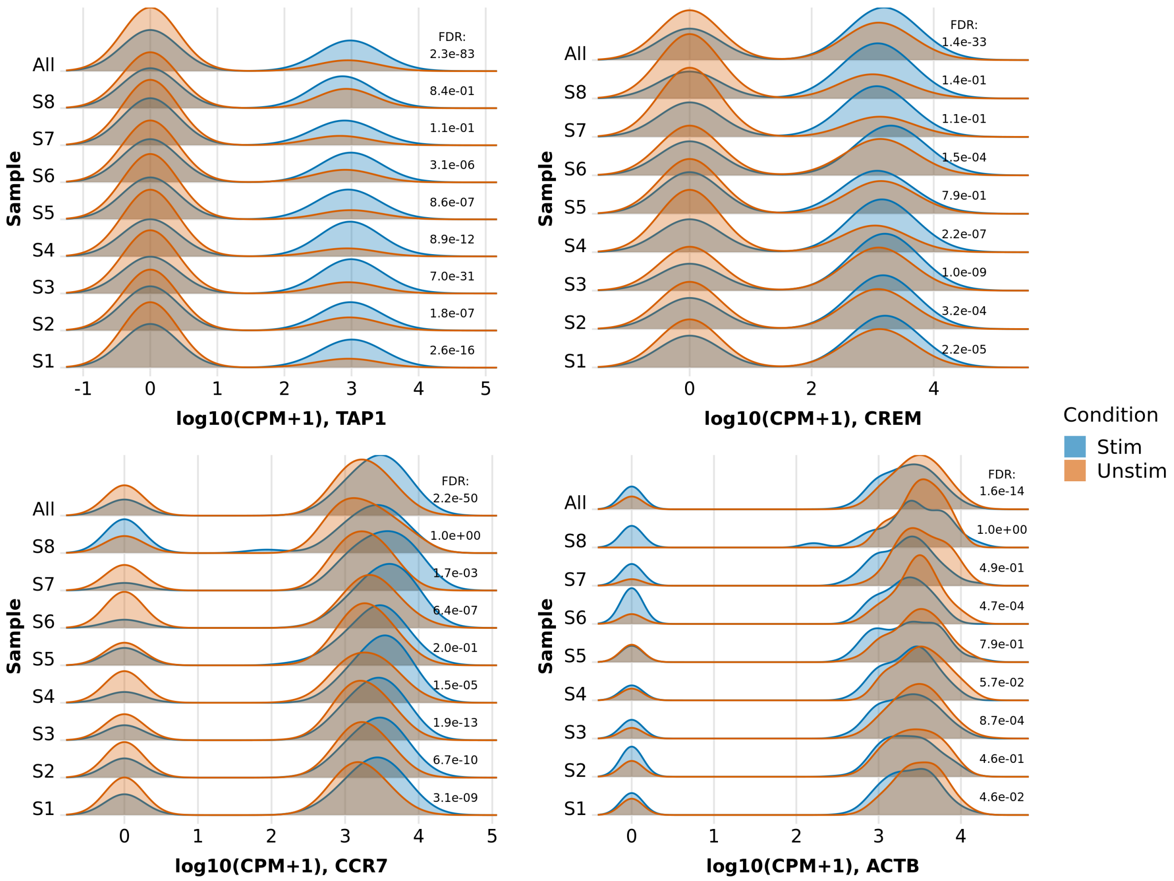


Figure S6: Example genes that are differentially expressed in B cells in part of the samples. The distributions of expression levels in both conditions are shown, either considering each sample separately or considering aggregated data. FDR: adjusted P value (Benjamini-Hochberg) of each DE test between conditions with likelihood-ratio test. CPM: count per million.

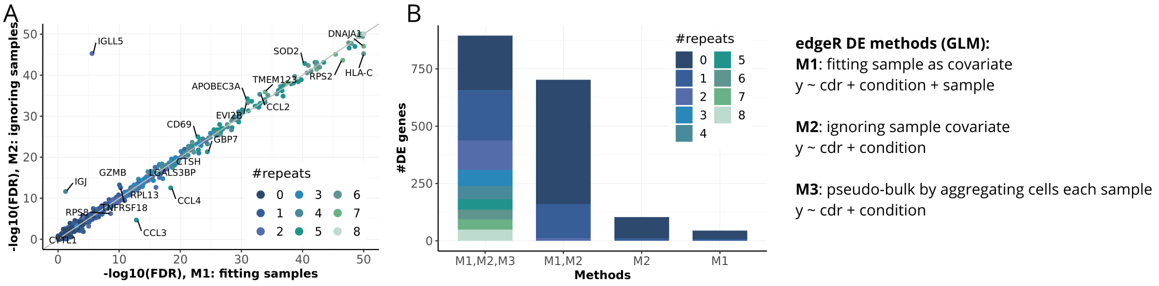


Figure S7: Differential expression analysis by aggregating multiple samples with three methods. Method 1 (M1): fitting sample as a covariate when performing DE analysis with treating each cell as a replicate; Method 2 (M2): ignoring sample covariate and treating each cell as a replicate; Method 3 (M3): summarising cells in each sample and treating each sample as a replicate, namely in a pseudo-bulk manner. (A) The scatter plot of -log10(FDR) between method 1 (M1) and method 2 (M2). Only genes with -log10(FDR) $\leq$50 are presented and genes with FDR differs $\geq$100 times between M1 and M2 are labeled. (B) The number of DE genes detected by different methods combinations, which again are categorized by the repeating times among the 8 samples. The notation for the three different DE analysis strategies is shared for (A) and (B).

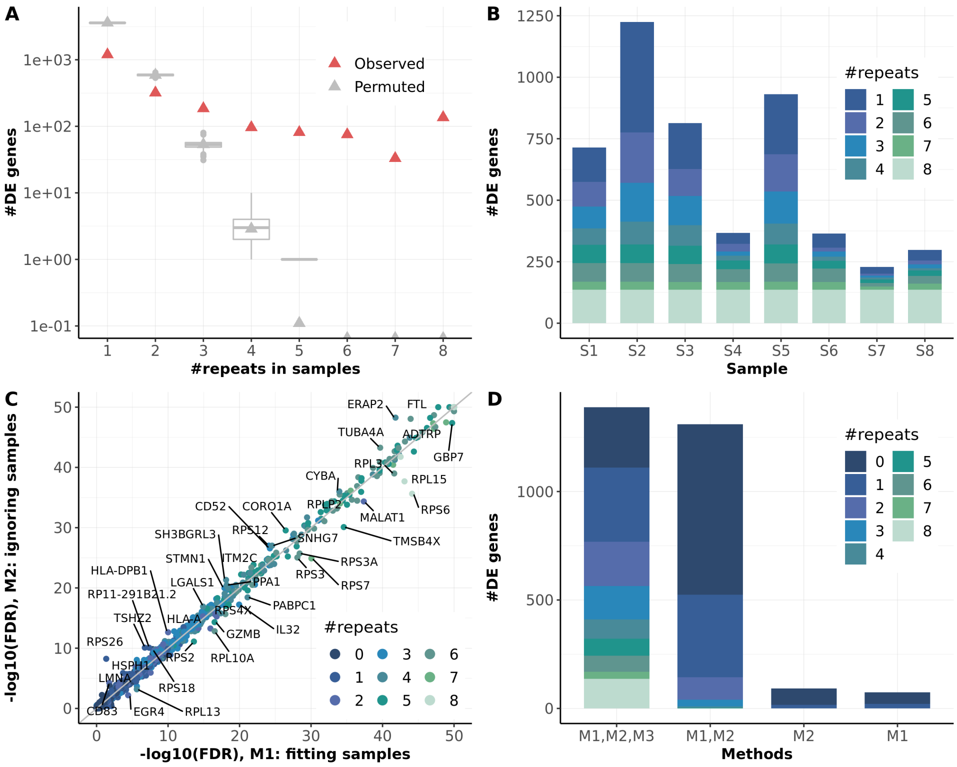


Figure S8: Differential expression analysis on CD4 T cells. (A) The number of repeating DE genes between stimulation and control, detected in one to eight individuals. The box plots in gray denote the results expected by chance (using 200 permutations). (B) The number of DE genes in each of the 8 samples, which are categorized by the repeating times, that is the number of individuals in which the gene is found to be DE. (C) The scatter plot of -log10(FDR) between method 1 (M1) with fitting sample as a covariate and method 2 (M2) with ignoring sample covariate. Only genes with -log10(FDR) $\leq 50$ are presented and genes with FDR differs $\geq 100$ times between M1 and M2 are labelled. (D) The number of DE genes detected by different methods combinations, which again are categorized by the repeating times among the 8 samples. M3 is the DE analysis in a pseudo-bulk manner. The notation for the three different DE analysis strategies is shared for (C) and (D).

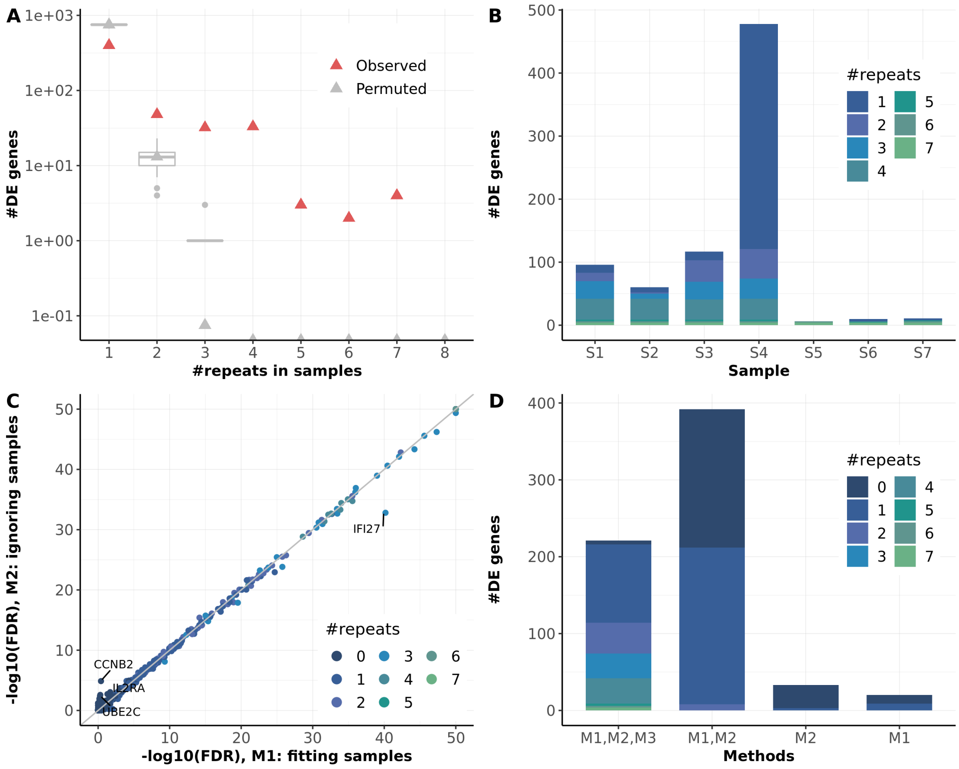


Figure S9: Differential expression analysis on CD8 T cells. The figure format is the same as Supp Fig S8.

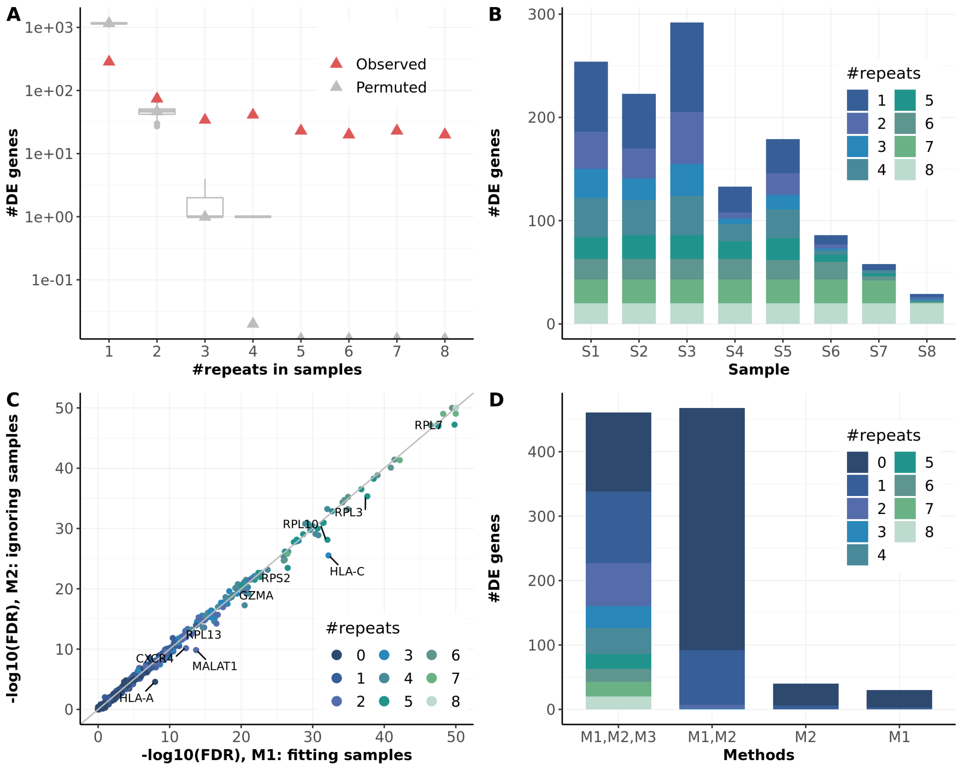


Figure S10: Differential expression analysis on NK cells. The figure format is the same as Supp Fig S8.

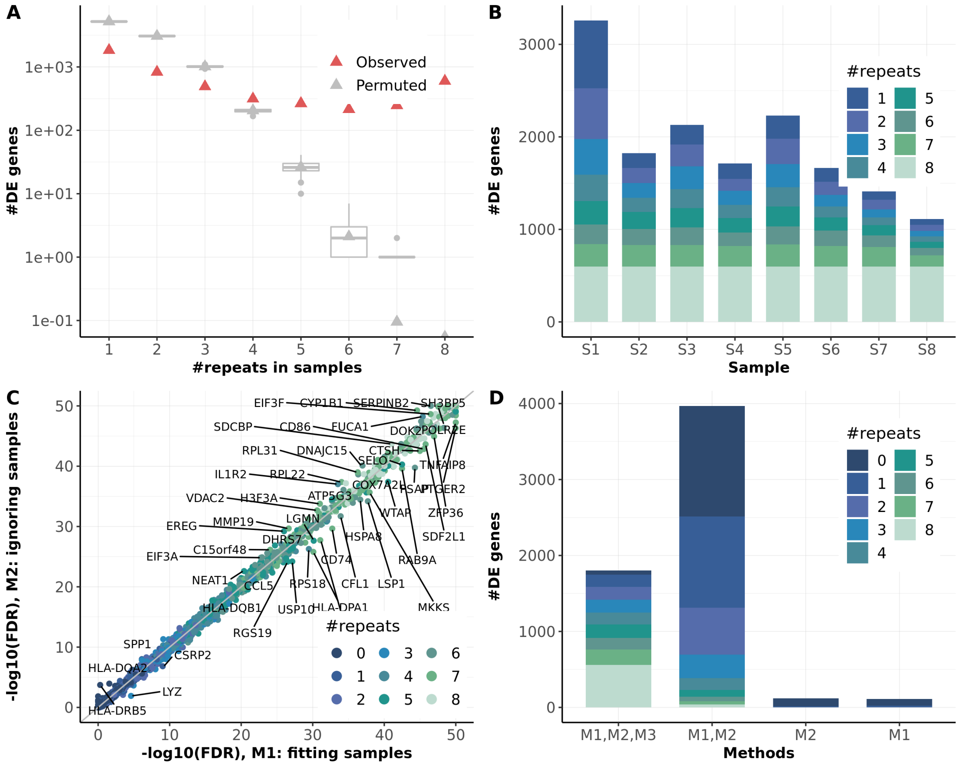


Figure S11: Differential expression analysis on CD14+ monocytes. The figure format is the same as Supp Fig S8.

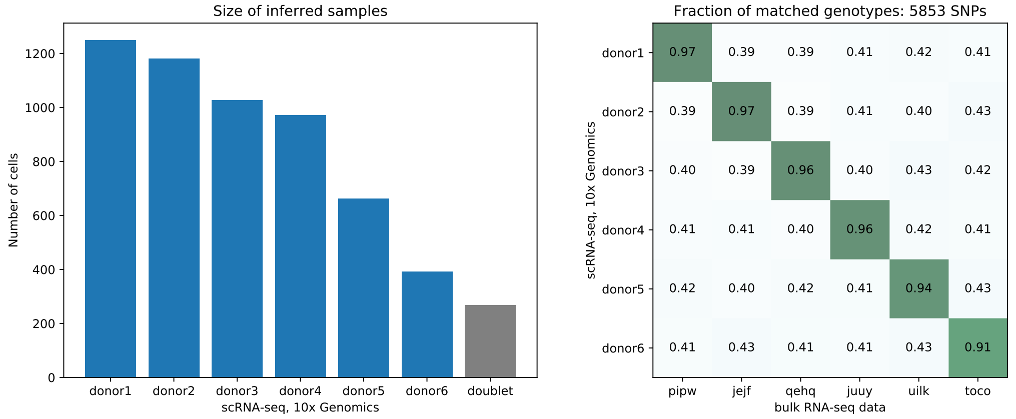


Figure S12: Sample alignment between pooled scRNA-seq data and bulk RNA-seq data from HipSci Project (see Methods). Left panel: size of inferred samples from a six-sample pooled scRNA-seq data based on 10x Genomics platform. Right panel: alignment between bulk RNA-seq samples to inferred samples from scRNA-seq data with Vireo by comparing the estimated genotype. The value of heatmap is the fraction SNPs with matched genotype between single-cell and bulk RNA-seq samples.

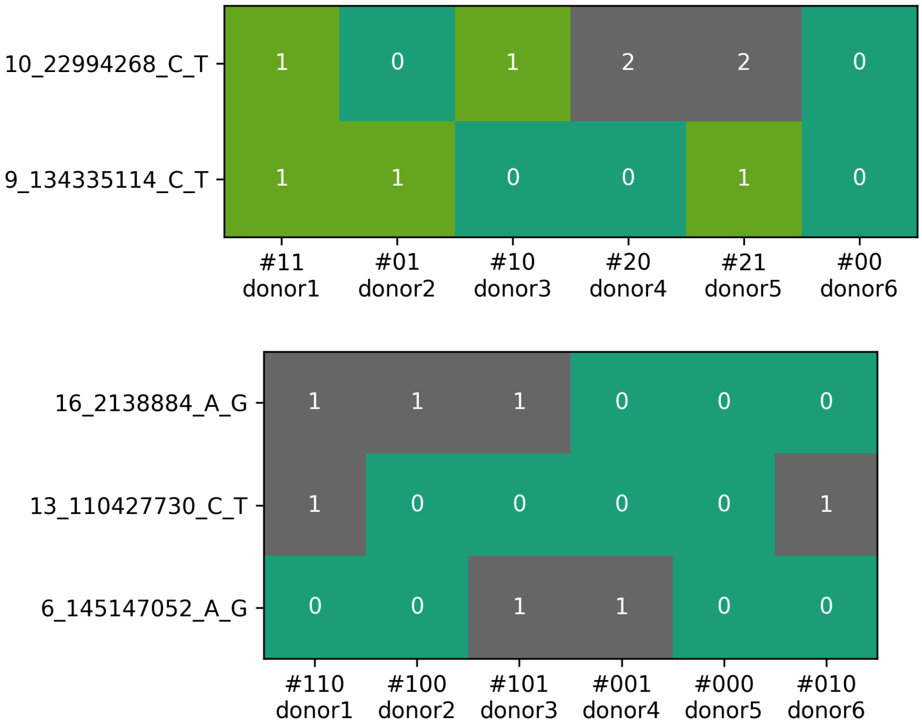


Figure S13: Identification of discriminatory variants from estimated genotype for the six pooled samples as in Supp. Fig. S12. Upper panel: two variants with three genotype categories. 0: homozygous reference allele; 1: heterozygous alleles; 2: homozygous alternative allele. Bottom panel: three variants without homozygous alternative allele.
